# Ectodysplasin overexpression reveals spatiotemporally dynamic tooth formation competency in stickleback and zebrafish

**DOI:** 10.1101/2025.05.01.651241

**Authors:** Zoe Z. Chen, Sujanya N. Narayanan, Lena M. Stagliano, Peter Q. Huynh, Shivani Sundaram, Emma J. Mackey, Craig T. Miller, Tyler A. Square

**Affiliations:** Department of Molecular & Cell Biology, University of California, Berkeley, CA 94720, USA; Department of Microbiology and Cell Science, University of Florida, Gainesville, FL 32611, USA

## Abstract

Organ initiation is often driven by extracellular signals that activate precursor cells competent to receive and respond to the signal, yet little is known about how dynamic competency is in space and time during development. Teeth are excellent organs to study organ initiation competency because they can be activated with the addition of a single signaling ligand, Ectodysplasin (Eda). Eda, a Tumor Necrosis Factor (TNF) ligand, is a critical regulator of ectodermal organ development, including teeth, acting through TNF receptors, like Edar, to activate NF-κB signaling in tooth precursor cells. While Eda is both necessary for normal tooth formation and sufficient for ectopic tooth formation, the spatial and temporal dynamics of competency for ectopic tooth initiation and maintenance remains unknown. To investigate the role of Eda in tooth specification, we generated transgenic sticklebacks and zebrafish with heat shock-inducible *Eda* overexpression. We find that stickleback *Eda* can drive de novo tooth formation in at least eight distinct oropharyngeal and cranial domains. Both zebrafish and stickleback exhibit maximal responsiveness to *Eda* overexpression during the critical window of pioneer tooth initiation, highlighting that the precursors of ectopic pharyngeal teeth likely follow a similar developmental trajectory as endogenous tooth precursors. Furthermore, we observe that some induced dental fields often undergo tooth regeneration to maintain themselves, allowing them to persist for months after the cessation of transgene activation. Finally, analysis of TNF receptor expression in sticklebacks reveals that ectopic tooth formation in the pharynx correlates with *Edar* and *Troy* expression, while in the region where teeth can form on the face, only *Troy* expression was detected, providing a possible molecular mechanism of competency involving spatially restricted receptor expression. These findings underscore the latent developmental potential, i.e. competency, of the vertebrate dentition and provide insights into organ competency during embryonic and post-embryonic development.

## Introduction

The localization of organ initiation is a highly regulated process that ultimately shapes a given body plan. Many organs are specified by secreted signaling ligands, which allows for some coordination between tissues during organ initiation (Perrimon et al., 2012). For cells to demonstrate “competency” to respond to a given signal, they must present a proper complement of receptors and other signal transduction machinery to respond to said ligand by specifying an organ. The extent to which cells are competent to initiate the formation of a given organ type, and how this competency changes throughout development, remains largely unknown. In vertebrates, internal organs typically exist as single units or in pairs that are specified with a conserved connectivity to other organs and tissues. Conversely, epithelial organs like teeth, hair, feathers, and scales tend to be more labile in their position and number both within and between species. Epithelial organs thus demonstrate a unique window into understanding how the morphological location of organ initiation can be altered, because the vertebrate body plan has withstood such variation while maintaining viability.

Teeth can vary especially widely in morphological location between species (Berkovitz and Shellis, 2023a). Similarly, the overall number of teeth present in a given tooth field can also vary at homologous morphological locations between species, or sometimes even within species (Ellis et al., 2015; Miller et al., 2014; Oeschger et al., 2022). Meanwhile, the cell types that constitute tooth organs demonstrate strikingly conserved developmental genetic programs across vertebrates (Gruenhagen et al., 2022; Rasch et al., 2016; Rostampour et al., 2019). Dental mesenchyme and epithelium coordinate through various extracellular signaling pathways to differentiate into odontoblasts and ameloblasts, respectively, which secrete bony tissues known as dentine and enamel (or enameloid), respectively. Taken together, these characteristics of tooth development represent an interesting case study in morphological evolution: conserved tooth units are specified using genetic cascades that must be localized in a comparatively plastic fashion, allowing teeth to “jump” around body plans with relative ease both experimentally and over evolutionary time. In particular, ray-finned fishes have accrued an impressively diverse array of dental morphologies. Across this group, at least 16 different oropharyngeal head skeleton elements naturally sport teeth in at least one lineage (Stock, 2007; Vandewalle et al., 2000), while a few species of fish also specify small teeth externally (known as denticles) (Greenwood and Greenwood, 1968; Mori and Nakamura, 2022).

Ectodysplasin (Eda) is a Tumor Necrosis Factor (TNF) which is known to bind at least one class of Tumor Necrosis Factor Receptor (TNFR), Ectodysplasin Receptor (Edar) (Elomaa et al., 2001). Once bound by Eda, Edar activates the Nuclear Factor Kappa B (NF-κB) pathway which induces transcription of genes involved in epithelial organ specification, including teeth (Liang et al., 2019; Ohazama and Sharpe, 2004; Schmidt-Ullrich et al., 2006). Eda has been shown to be both a necessary factor for proper dental development, as well as sufficient to drive the differentiation of ectopic and supernumerary teeth. Human *EDA* mutants present a disease known as Ectodermal Dysplasia (for which *EDA* was named), typically characterized by dental abnormalities ranging from one or more missing teeth (tooth agenesis) to complete loss of teeth (edentulousness) (Liu et al., 2022; Zeng et al., 2017). Fish models, including sticklebacks and zebrafish, also demonstrate stark losses of teeth when *Eda* is mutated, resulting in significantly reduced tooth numbers (Harris et al., 2008; Wucherpfennig et al., 2019). Human clinical trials have shown that prenatal exposure to a recombinant human EDA (Fc-EDA) is associated with a less-severe reduction in primary tooth number than in untreated siblings with the same *EDA* genotype (Schneider et al., 2018). Previous overexpression experiments utilized transgenes that drove the zebrafish *eda* gene under the control of the ubiquitous *ef1a* promoter in zebrafish and *Astyanax* models, finding that both species were capable of differentiating ectopic teeth (Aigler et al., 2014; Jandzik and Stock, 2021). Provocatively, the ectopic teeth that arose in these experiments were often morphologically discontiguous with endogenous tooth fields, meaning they did not represent simple expansions of existing tooth fields, but rather represented truly *de novo* deployment of a tooth differentiation program at distinct head skeleton elements. Furthermore, the ectopic tooth fields that arose in these experiments occur naturally in other fish species, suggesting that these experimental phenotypes phenocopy existing natural variation across fish groups. Notably, some of the experimentally induced teeth in *Astyanax* were shown to have been repeatedly lost and regained in parallel within their order (Characiformes). This response to exogenous Eda suggests that a latent developmental potential to form teeth has persisted for 100 million years or more, having been reactivated in separate lineages multiple times in order to reinvoke teeth at certain oropharyngeal bones (Aigler et al., 2014; Jandzik and Stock, 2021). However, these previous studies used ubiquitous overexpression of *eda*, and thus lacked temporal control of *Eda* overexpression, raising the question of how temporally labile tooth formation competency is during embryonic and post-embryonic development.

To test the spatial and temporal windows of tooth formation competency, we developed transgenic lines of sticklebacks and zebrafish capable of heat shock-inducible *Eda* expression. Our transgenes use the zebrafish *hsp70l* promoter to overexpress an *mCherry-P2A*-*Eda* polycistronic gene in each zebrafish and stickleback, where the *Eda* coding region supplied to each species encodes their endogenous protein. Overall, these transgenic fish lines allow us to test (1) whether competency is spatially restricted in two different species (2) whether competency is temporally dynamic, i.e. whether some regions are competent to respond during some developmental windows but not others, and (3) whether ectopic teeth are retained or are capable of regeneration after exogenous *Eda* overexpression is discontinued. We find that *Eda* overexpression indeed elicits ectopic tooth formation in sticklebacks, which can specify teeth in at least eight unique dental domains, while zebrafish are limited to the previously reported single (paired) ectopic tooth domain in the dorsal pharynx. We further find that zebrafish and sticklebacks both demonstrate the highest potential to form ectopic pharyngeal teeth during the window wherein their endogenous pioneer teeth are specified. We also observed evidence of tooth regeneration at ectopic tooth fields multiple months after heat shock activation of *Eda* during embryonic stages. Finally, cross-referencing ectopic tooth formation profiles with TNFR gene expression in sticklebacks provides evidence that *Edar* and *Troy*, but not *Relt*, are expressed in regions coincident with ectopic tooth formation competency during periods where ectopic teeth are most receptive to initiation, providing a possible mechanism involving receptor expression for the dynamic spatial and temporal competency.

## Results

### *Eda* gene duplication and differential paralog retention

Some fish genomic assemblies house more than one annotated *Eda* gene. A previous synteny analysis found that some teleost fish species possess two copies of *Eda*, while the spotted gar, a non-teleost, possesses only one copy, suggesting that these duplicates arose via the teleost whole-genome duplication (TGD) (Braasch et al., 2009). To further understand the evolutionary relationships among *Eda* loci in jawed vertebrates, we generated a phylogeny using Eda amino acid sequences from a range of diverse fish genomes (Fig. S1). Overall, this phylogeny provides additional evidence that *Eda* gene duplicates indeed originate from the TGD. After duplication, both resulting *Eda* paralogs were retained by many, but not all teleost lineages, as each paralog group formed separate clades that generally recapitulate widely-accepted relationships among major fish groups (Betancur-R et al., 2017). As previously hypothesized (Braasch et al., 2009), our amino acid similarity analysis supports a scenario where zebrafish (and other Cypriniforms) have retained the opposite *Eda* paralog as sticklebacks (and other Neoteleosts); following the previous designations, here we refer to the paralog retained by sticklebacks as the “A” paralog and the paralog retained by zebrafish as the “B” paralog. Additionally, some groups appear to have further duplicated and/or lost one *Eda* paralog: two B group *Eda* genes were detected in two separate clades, carp+goldfish and salmonids. The two species of salmonids assessed here thus appear to retain three *Eda* genes, one A and two B paralogs, while carp and goldfish have two *Eda* genes, both of which are B paralogs.

### *Eda*-driven ectopic tooth formation competency in stickleback can be triggered during or after endogenous tooth specification

To test whether a type A *Eda* can cause ectopic tooth development in sticklebacks, we generated a stickleback cross using the *Eda* OE transgene and a *dlx2b:eGFP* reporter transgene (Jackman et al., 2022; Jackman and Stock, 2006). This zebrafish *dlx2b* reporter transgene is expressed in a tooth-specific manner in sticklebacks, as in zebrafish, allowing us to detect nascent tooth germs in living fish. We split this cross into two groups for two non-overlapping overexpression (OE) treatments to test whether ectopic tooth specification required exogenous Eda during the same window wherein endogenous teeth were specified. In the first half of the cross, we overexpressed stickleback *Eda* during embryonic stages from 1-9 days post fertilization (dpf) by administering ∼1 hour heat shocks (see methods) every 12 hours (16 total heat shocks) followed by a six-day recovery period (Fig. 1A). This treatment encompassed the establishment of all endogenous tooth fields, spanning from early neural crest cell migration to pioneer tooth differentiation. We found that 100% (n=8/8) of GFP+, *Eda* OE sticklebacks grew multiple ectopic teeth, whereas 0/7 control (mock heat shocked, non-OE fish) and 0/8 non-heat shocked, GFP+ OE transgene carriers demonstrated tooth organs outside of their described endogenous tooth fields (Fig. 1B-D, brackets). All heat shocked *Eda* OE fish demonstrated a higher number of oral tooth germs, and in 5/8 individuals the dentary tooth organs were arranged in a cluster as opposed to the single row seen in the control individuals (Fig. 1E-H, brackets). In the pharynx, multiple ectopic teeth formed along the ventral midline (Fig. 1I-L). At ceratobranchial 4 (cb4) bones, clusters of 3-6 ectopic teeth were detected in 8/8 individuals. At hypobranchial 3 (hb3), clusters of 2-4 ectopic teeth were observed in 8/8 fish, at hb2 we observed 1-2 ectopic teeth in 3/8 fish, and at hb1 we observed 1-2 ectopic teeth in 7/8 fish. We additionally detected basihyal (bh) tooth organs in 2/8 individuals (Fig. 1F, arrow). At cb5, Alizarin Red staining revealed additional supporting bone (bone of attachment) at cb4 and cb5 in 8/8 fish, sometimes in regions without ectopic teeth (Fig. L, black arrow). In 3/8 fish, the left and right endogenous cb5 pharyngeal tooth fields were fused across the midline (Fig. 1L, arrowhead).

**Figure 1.**
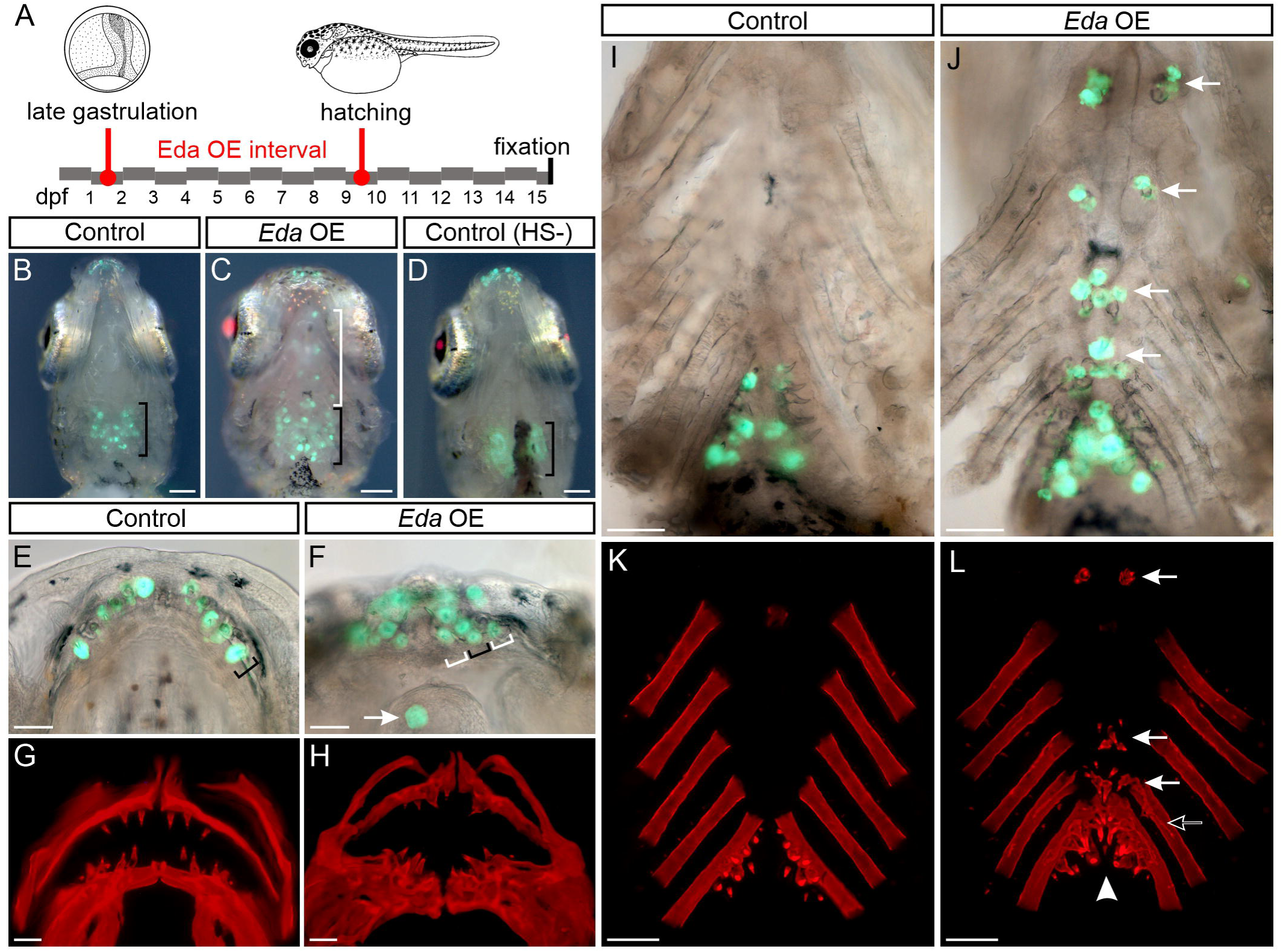
Embryonic *Eda* overexpression in sticklebacks. (A) an outline of the embryonic treatment. Fish were heat shocked from 1 to 9 dpf 2x per day, from late gastrulation to hatching. Drawings modified from Swarup, 1958. (B-D) Ventral views (anterior to top) of fish carrying the *dlx2b*:*eGFP* reporter, marking nascent tooth germs (green). A black bracket marks the anterior-posterior extent of endogenous pharyngeal tooth germs in B-D. White bracket in C indicates ectopic tooth germs. B shows a heat shocked control fish that does not carry the *Eda* overexpression (OE) transgene. C shows a heat shocked treatment fish that does carry the *Eda* OE transgene (mCherry). D shows an un-heat shocked control fish that carried the *Eda* OE transgene (mCherry in lens only). The individuals in B and C are imaged after heart removal, during pharyngeal extraction; the individual in D is a future treatment fish for the experiment depicted by Fig. 2, and is thus imaged under light anesthesia without heart removal. Note the magenta mCherry signal in panels C and D relative to B. In panel D, the green and red channel fluorescence gain settings were higher than in B,C to highlight the pharyngeal tooth distribution and red lens. (E,F) mandibular tooth germs marked by the *dlx2b:eGFP* transgene in control (E) and *Eda* OE fish (F). Black brackets in E,F indicate a single row of teeth, white brackets in F indicate supernumerary tooth rows. Arrow in F indicates a tooth on the anterior basihyal (bh). (G,H) Alizarin Red oral skeleton preparations from control (G) and *Eda* OE (H) individuals. (I-J) ventral pharyngeal tooth germs marked by the *dlx2b* transgene in control (I) and *Eda* OE fish (J). White arrows in J mark ectopic tooth germs. (K,L) Alizarin Red ventral pharyngeal skeleton preparations from control (K) and *Eda* OE (L) fish. Arrows in L mark ectopic teeth. Black arrow in L marks ectopic bone of attachment. Arrowhead in L indicates an instance of induced pharyngognathy, i.e midline fusion of ventral tooth fields. Scale bars in B-D=200 μm; E-H=50 μm; I-L=100 μm.

To test whether ectopic teeth could be activated at developmental stages after neural crest cell migration and the initiation of endogenous pioneer tooth differentiation, we overexpressed *Eda* during larval stages in the remaining individuals of the cross described above. We administered heat shocks once every 12 hours from 17-25 dpf (16 total heat shocks), again followed by a six-day recovery period (Fig 2A), i.e. the same duration and cadence of heat shocks and the same recovery period as administered to the first group but shifted to a later developmental stage. This treatment interval corresponds to late larval feeding and growth stages, after each endogenous tooth field has been established. We assessed GFP fluorescence in those fish carrying the *dlx2b:eGFP* reporter, and later stained all individuals with Alizarin Red. We found that only 3/23 fish formed ectopic pharyngeal teeth resulting from this treatment, where 2/23 demonstrated 1-2 cb4 teeth, and 1/23 demonstrated hb1 and bh teeth (Fig. S2). This result suggests that the potential to form ectopic pharyngeal teeth upon exposure to exogenous *Eda* is still maintained, albeit at a lower efficacy, long after the bulk of neural crest cell migration and endogenous tooth field establishment. Interestingly, a distinct class of ectopic teeth was additionally prompted by this treatment: we found that 20/23 of these *Eda* OE fish specified tooth organs on their face (Fig. 2B-E). These face teeth were most typically located superficially to the anteriormost two infraorbitals (io1 and/or io2), as found in 20/23 *Eda* OE fish (Fig. 2C, arrows). Face teeth were also observed on the ventrolateral sides of the dentary, articular, and/or angular bones in 17/23 *Eda* OE fish (Fig. 2C, arrowhead). Alizarin Red staining revealed a general lack of bone of attachment surrounding the bases of most face teeth, unlike the observed effect in the pharynx (Fig. 2E). Notably, when bone of attachment was observed, it was always on or near the ventral dentary.

**Figure 2.**
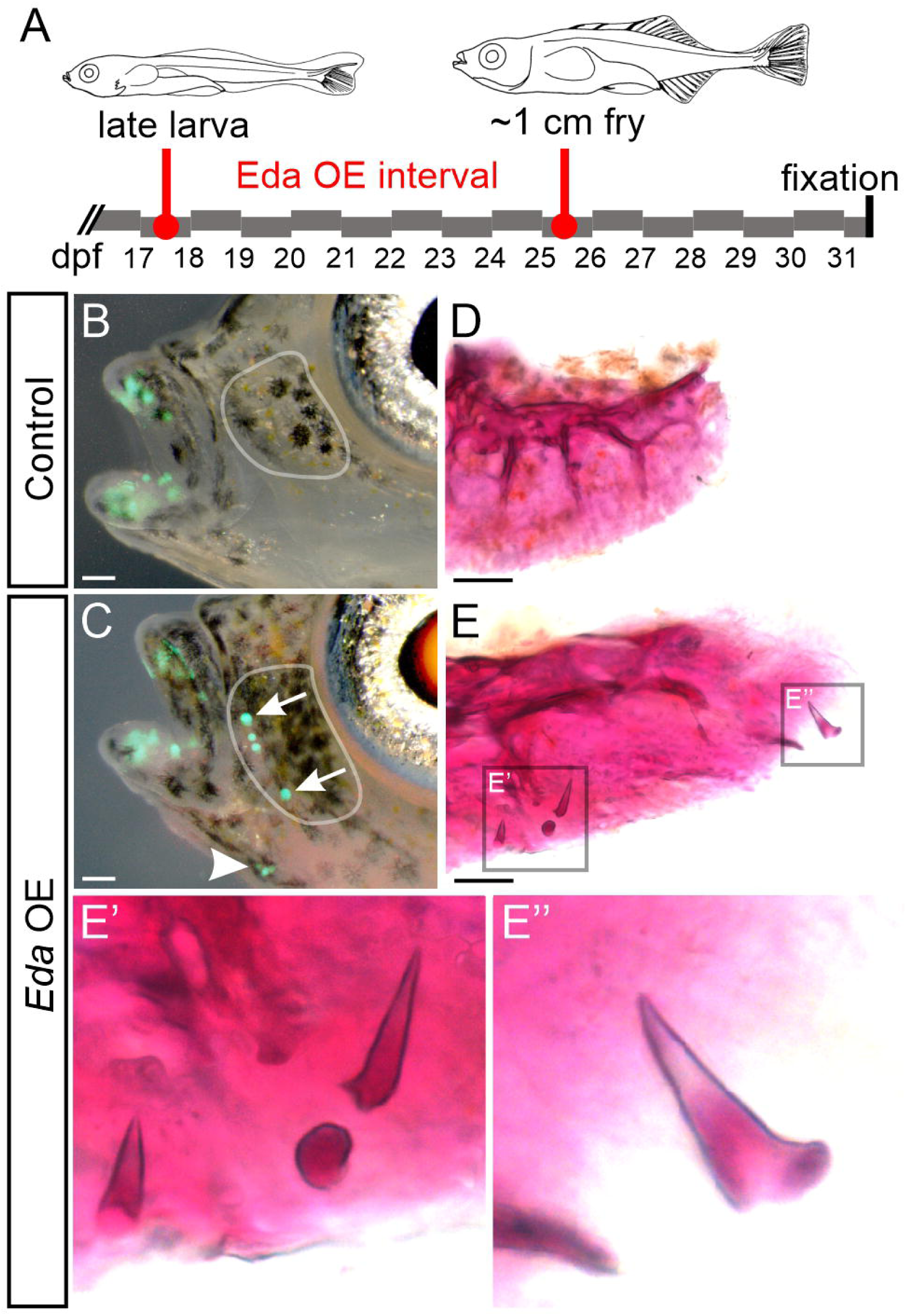
Larval *Eda* overexpression in sticklebacks. (A) an outline of the late larval treatment. Fish were heat shocked from 17 to 25 dpf 2x per day. Drawings modified from Swarup, 1958. (B,C) Larval *Eda* OE causes face tooth germ initiation in *Eda* OE (C) but not control fish (B). Arrows in C mark the ventralmost and dorsalmost infraorbital (io) teeth, arrowhead marks a cluster of ventral dentary (v.dent) teeth. Gray outlines indicate the position of io1, which is shown dissected and stained in panels D,E. (D,E) dissection of the first infraorbital from an *Eda* OE fish (E) reveals ossified teeth on the face, which were not present in control fish (D). Insets indicated in E are shown enlarged in E’ and E”. Scale bars in B,C=100 μm; D,E=50 μm.

### Tooth differentiation on the stickleback face is negatively correlated with neuromast differentiation

To test whether face tooth differentiation was spatially biased, we performed another late larval treatment by administering two heat shocks per day to overexpress *Eda* from 20-36 dpf (32 total heat shocks). By counterstaining with DASPEI, which marks neuromasts (yellow in Fig. 3A,B), we first documented the spatial distribution of face teeth by using neuromast positions as reference points (Fig. 3C). We also counted a subset of neuromasts (those that form nearest the ectopic teeth, referred to hereafter as “anterior neuromasts”) to test whether neuromast number demonstrated a relationship with facial tooth number. DASPEI staining alongside the *dlx2b:eGFP* reporter revealed that these facial tooth organs were always situated near neuromasts (Fig. 3B’, arrowheads), and occasionally they appeared to form in place of neuromasts (Fig. 3B’, arrow).

**Figure 3.**
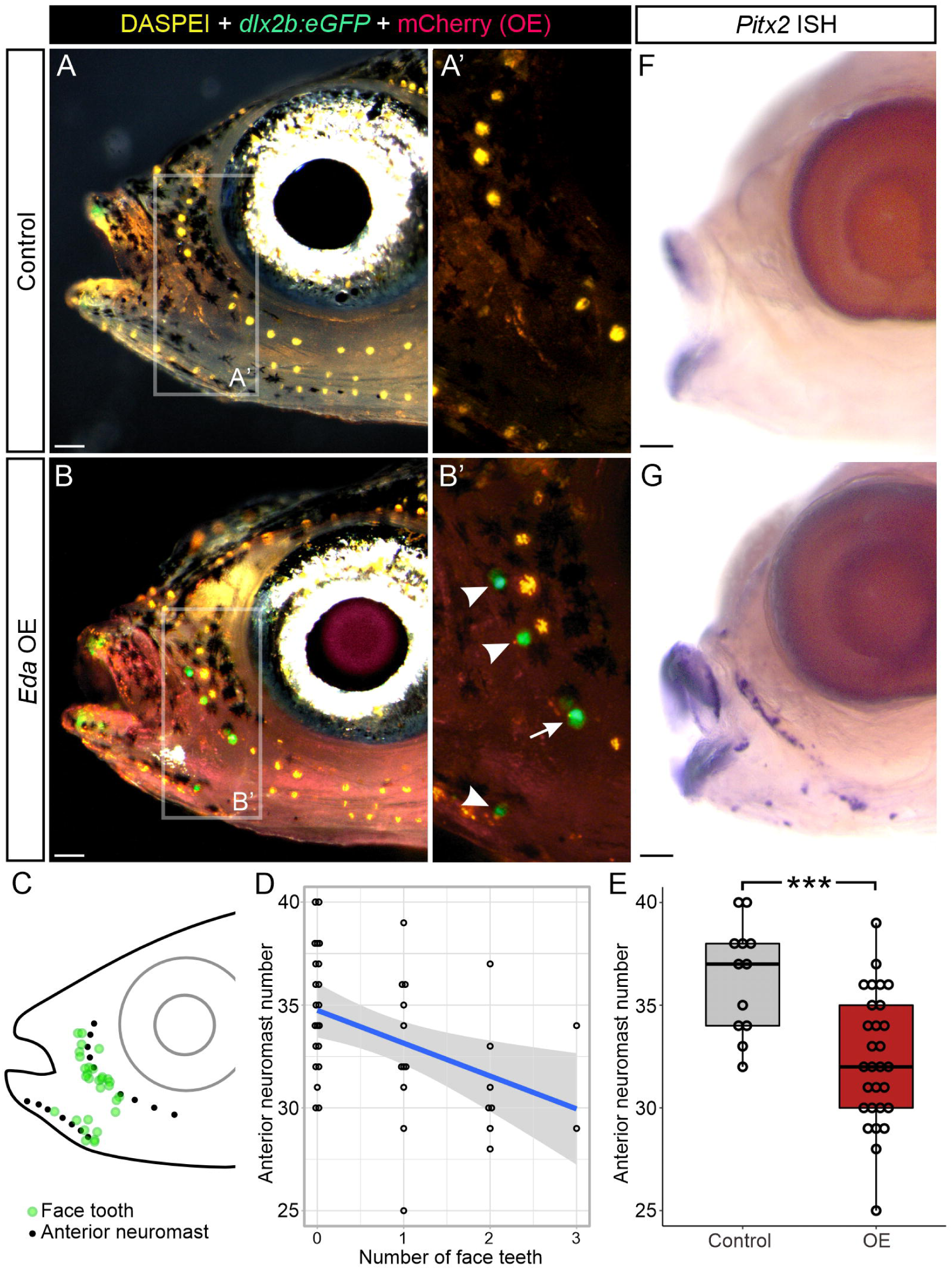
Face tooth location and negative relationship with anterior neuromasts in sticklebacks. (A,B) Left lateral views of control (A) and *Eda* OE (B) fish that underwent a larval heat shock treatment from 20-36 dpf. DASPEI shown in yellow, eGFP from the *dlx2b* tooth reporter shown in green, mCherry shown in magenta. A’ and B’ show the insets indicated in A and B, sans the brightfield overlay. Arrowheads in B’ mark ectopic face teeth growing nearby neuromasts, while the arrow in B’ marks a face tooth growing in the position of a neuromast. (C) a diagram showing neuromast (black dot) position relative to face tooth (green dot) position at 36 dpf. (D) A scatterplot of anterior neuromasts (pictured as black dots in panel C) as a function of the number of face teeth. A statistically significant negative linear correlation was observed (*P=*0.006). (E) A box and whisker plot showing a significant reduction in anterior neuromast counts for *Eda* OE vs. the control condition (mock heat shock). Wilcoxon Rank Sum test *P=*0.001 (***) n=12 control vs n=29 *Eda* OE fish. (F,G) *Pitx2 in situ* hybridization (ISH) revealed ectopic expression coincident with the region where ectopic teeth form in the *Eda* OE condition but not the control condition. Scale bars=100 μm.

We reasoned that if face teeth sometimes arose from repurposed neuromast progenitors, we would expect to see a negative correlation between anterior neuromast number and the presence of face teeth. Consistent with our observations that some face teeth appear to form in place of certain neuromasts, we found a significant negative linear relationship between the number of face teeth and the number of anterior neuromasts observed in each individual fish (*P*=0.006; Fig. 3D). Overall anterior neuromast number was also significantly reduced in the *Eda* OE condition (*P*=0.001; Fig. 3E). These results are consistent with the hypothesis that the neuromast cell linage donates cells to face tooth formation, though notably there are other possible explanations for this negative correlation, e.g. that Eda negatively affects neuromast differentiation directly and separately from its role in promoting tooth differentiation. *In situ* hybridization for *Pitx2* on this same treatment condition revealed ectopic expression of this transcription factor in these same regions (Fig. 3F,G), suggesting that indeed these are *bona fide* teeth/odontodes rather than generic ectopic bone that is coincidentally conical.

### *Eda* overexpression causes accelerated differentiation of endogenous tooth germs

To ask whether endogenous tooth organ development could be accelerated by overexpression of a type A *Eda*, we again overexpressed *Eda* in sticklebacks on a *dlx2b:eGFP* tooth reporter background. We administered heat shocks once per day from 2-9 dpf (8x heat shocks), preceding endogenous oral tooth differentiation, and checked daily for eGFP signal. We observed accelerated pioneer tooth differentiation in the oral jaws (Fig. 4A,B, arrowheads): all treatment animals (n=9/9) had four oral tooth germs at 9 dpf, whereas no control fish had any detectable GFP signal in their mouth at this time point (n=0/10). Notably, the four GFP+ tooth germs observed in this arrangement – a bilateral pair of pioneer tooth germs on each their premaxillary and dentary tooth fields – is reminiscent of normal expression of the *dlx2b* enhancer at 12 dpf in WT fish (Fig. 4C), suggesting that endogenous tooth development is accelerated by *Eda* OE.

**Figure 4.**
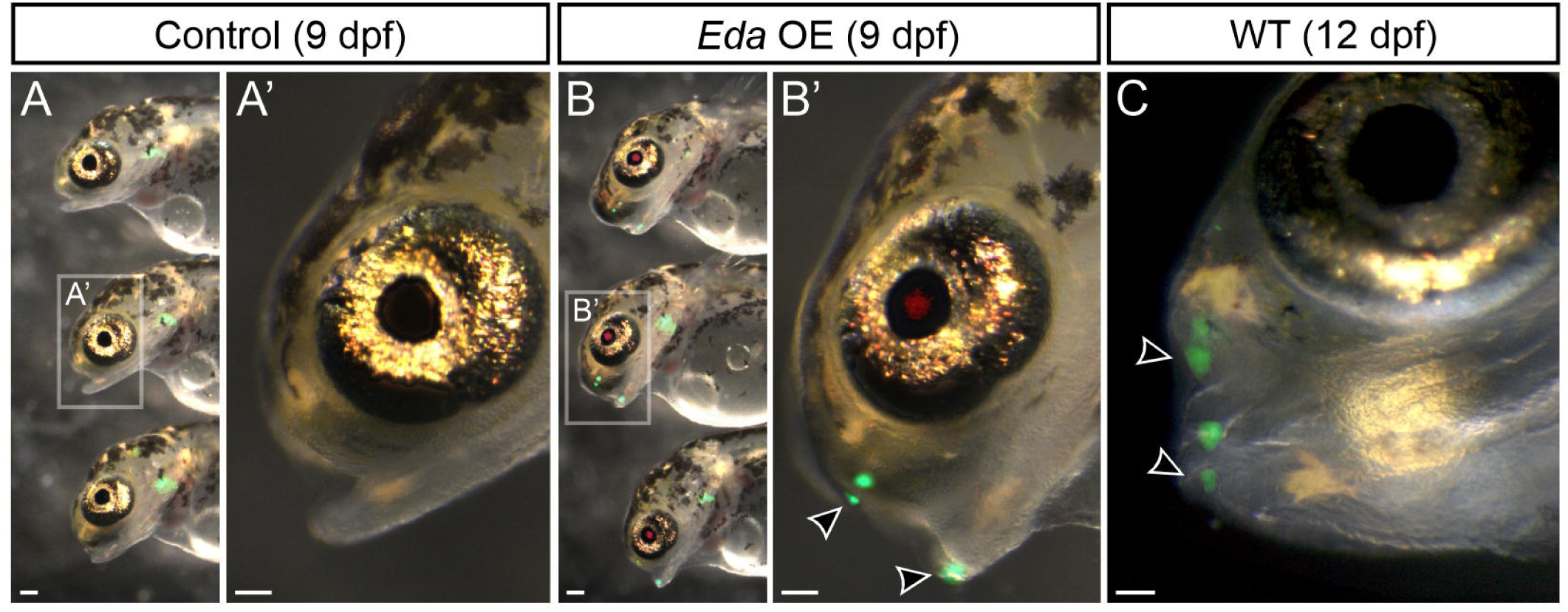
*Eda* overexpression causes precocious tooth differentiation in sticklebacks. (A,B) at 9 dpf, *Eda* OE fish demonstrated precocious oral tooth germs. Images show eGFP from the *dlx2b* tooth reporter and mCherry from the OE transgene. A’ and B’ show the fish indicated in A and B in greater detail. Black arrowheads indicate pioneer tooth germs. (C) at 12 dpf in WT, pairs of germs are observed on the premaxilla and dentary (black arrowheads), similar to the condition seen in the 9 dpf *Eda* OE fish, suggesting these represent accelerated endogenous teeth. Sale bars=100 μm.

### *Eda*-driven tooth formation competency in sticklebacks shows distinct dose-dependence and temporal requirements across different regions of the head

We next aimed to understand the anatomical and temporal windows of *Eda*-driven tooth formation competency in sticklebacks by further varying the timing and number of heat shock treatments (i.e. “doses” of *Eda*). We thus conducted a series of unique *Eda OE* treatments followed by an analysis of tooth ossification by Alizarin Red staining at 24 dpf for each treatment. See Fig. 5A for a summary of results of each treatment. These experiments revealed that not only are different skeletal elements “primed” for a tooth formation response during different developmental intervals, but also that different regions of the head skeleton demonstrate different propensities for ectopic tooth formation under a given heat shock regiment. For example, ectopic teeth were often observed anterior to the first epibranchial (eb1), positioned upon the medial-most gill raker that emanates from this element; these teeth displayed the widest interval during which they could be specified by a single heat shock, lasting from 4 to 10 dpf, compared to regions like hb1 and cb4 where the induction window was narrower, lasting from 5 to 7 or 8 dpf, respectively. Some regions appeared generally more recalcitrant to tooth induction via *Eda*, for example the basihyal (bh) never produced any teeth following a single heat shock, whereas multiple heat shocks could produce teeth in this region, though always at a lower rate compared to regions like cb4 or eb1 in the same treatment. Similarly, face teeth were only observed in multi-heat shock treatments, and the infraorbital teeth were always induced at a higher rate compared to those on the ventral dentary.

**Figure 5.**
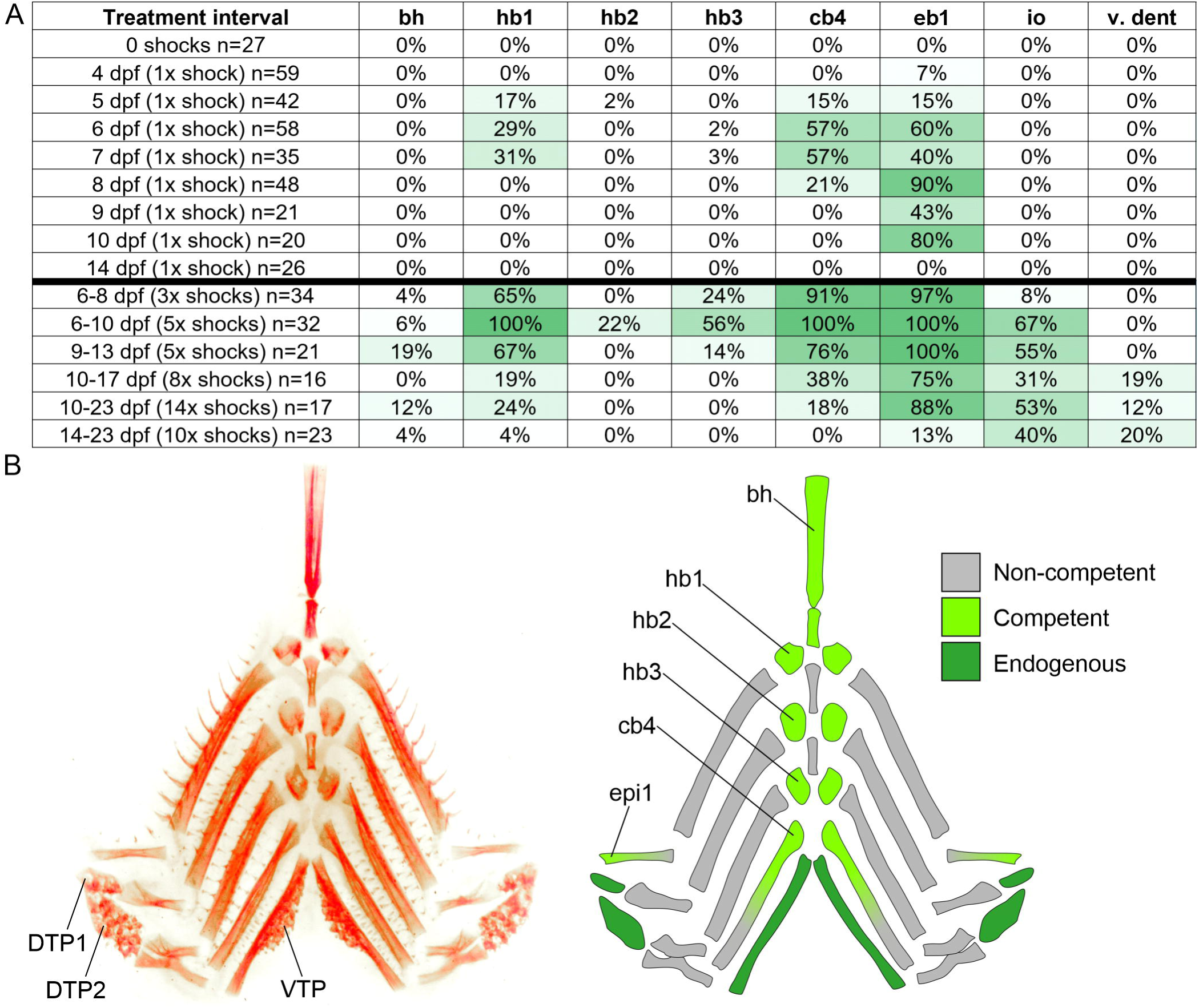
Stickleback tooth formation competency through ontogeny. (A) a heatmap showing the % occurrence of ectopic tooth formation at different morphological locations. The leftmost column describes each treatment, each other column represents a bone or morphological region where ectopic teeth were observed in the percentage of fish shown. Treatments below the black bar consisted of multiple heat shocks, those above are single heat shock experiments. Ectopic teeth were not detected in any negative control fish (n=0/169 mCherry-negative, heat shocked fish [not listed]; n=0/27 mCherry-positive, un-heat shocked fish). bh, basihyal; hb1-3, hypobranchial 1-3; cb4, ceratobranchial 4; eb1, epibranchial 1; io, infraorbital; v. dent, ventral dentary. (B) Alizarin Red-stained pharyngeal skeleton (left) and an illustration of the skeletal elements indicating whether each element ever bore teeth under at least one *Eda* OE treatment. Dorsal tooth plate 1 and 2, DTP1 and DTP2; ventral tooth plate, VTP. Other abbreviations same as panel A.

### *Eda*-driven tooth formation competency in zebrafish shows distinct dose-dependence and temporal requirements

To define the dose and temporal requirements for Eda in zebrafish ectopic tooth formation competency, we performed an array of heat shock experiments using the zebrafish *eda* coding sequence (a type “B” *Eda*) and analyzed ectopic dorsal pharyngeal tooth formation by Alizarin red at 10 dpf (Fig. 6A). Similar to sticklebacks, we found that the rates at which dorsal pharyngeal teeth formed in zebrafish upon *eda* OE were also dosage and time dependent (Fig. 6B). Zebrafish ectopic tooth formation competency also appears to peak surrounding endogenous pioneer tooth initiation, which occurs at ∼2 dpf (Van der Heyden and Huysseune, 2000): 24% of fish heat shocked at 2 dpf demonstrated at least one ectopic tooth. Conversely, only 6% and 3% of fish demonstrated at least one ectopic tooth when heat shocked at 1 or 3 dpf, respectively. Like in sticklebacks, multi-heat shock treatments usually demonstrated higher rate of ectopic tooth formation, to the exclusion of a double heat shock at 3 and 3.25 dpf.

**Figure 6.**
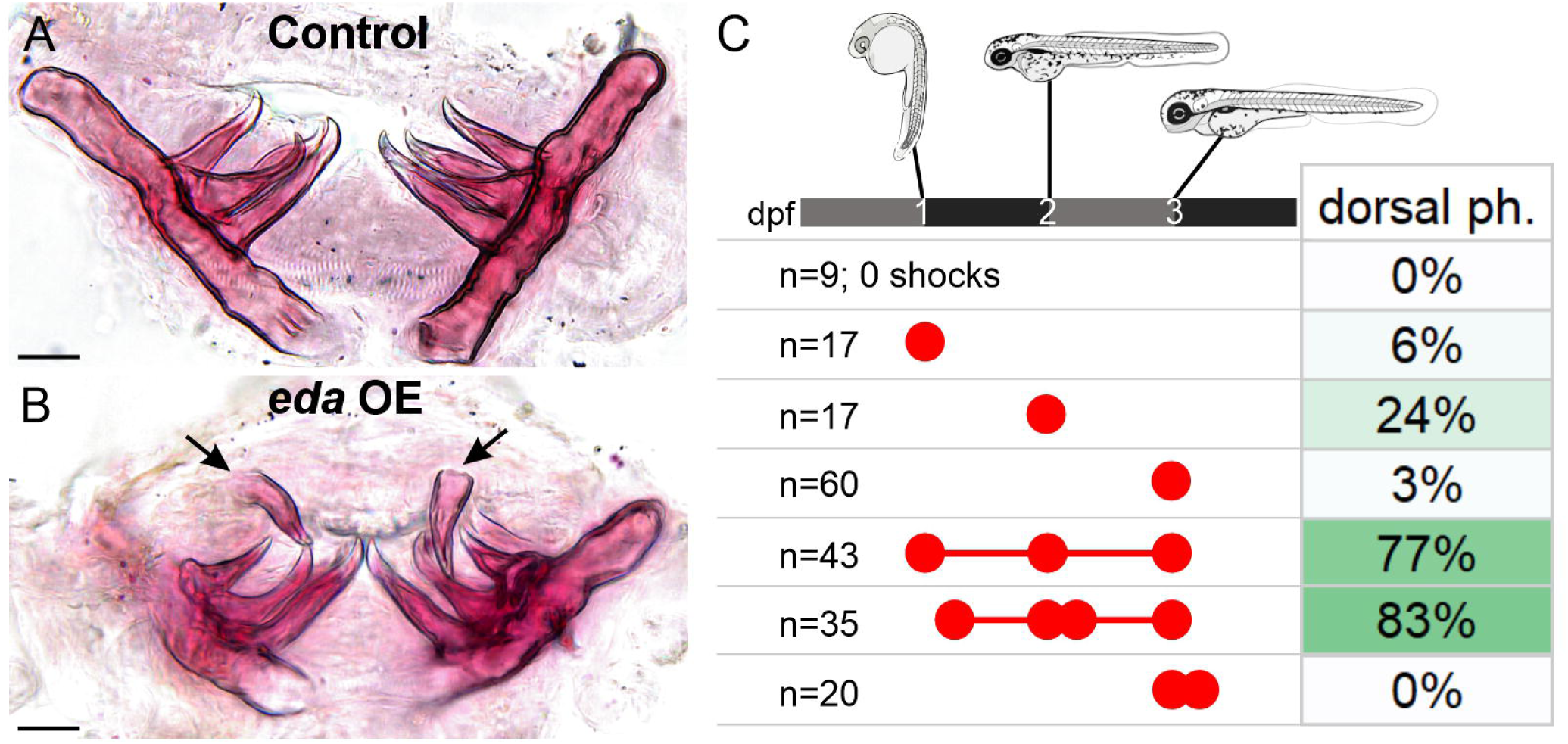
Zebrafish tooth formation competency though ontogeny. (A,B) Alizarin Red-stained pharyngeal tooth fields dissected from the 24+48+72h experiment showing a pair of bilateral ectopic teeth (arrows). Oral views, dorsal to top. (C) results of different heat shock treatments on ectopic tooth formation in zebrafish. Ectopic teeth were not detected in any negative control fish (n=0/36 mCherry-negative, heat shocked fish [not listed]; n=0/9 mCherry-positive, un-heat shocked fish). Red dots indicate the timing of heat shocks relative to the timeline shown at the top of C; treatments occurred at 24h, 30h, 48h, 54h, 72h, and/or 78h post-fertilization. Scale bars=25 μm.

### Ectopic pharyngeal but not face teeth are capable of regeneration without additional *Eda* overexpression

Previous *Eda* overexpression experiments in zebrafish and *Astyanax* utilized a *Xenopus Elongation Factor 1a* (*ef1a*) promoter that drives transcription in most zebrafish cell types at all life stages, including in adults (Amsterdam et al., 1996; Johnson and Krieg, 1994; Moon et al., 2013). Since this promoter remains active during adult stages, it remained unknown whether ectopic teeth could be maintained if *Eda* OE was discontinued. To test if ectopic teeth could be retained in zebrafish after the cessation of *Eda* OE, we overexpressed zebrafish *eda* at 24, 30, and 48hpf, followed by a two month recovery period, at which point we assayed upper pharyngeal histology by Hemotoxylin and Eosin staining on sections or by Alizarin Red wholemount staining (Fig. 7A,B). We found ectopic teeth were present in n=4/10 fish with the OE construct, and in every individual where ectopic teeth were observed, at least one of those teeth was an unerupted tooth germ associated with a nearby erupted tooth (Fig. 7B, arrows), suggesting that tooth replacement via regeneration was ongoing two months after the last dose of *Eda* was administered.

**Figure 7.**
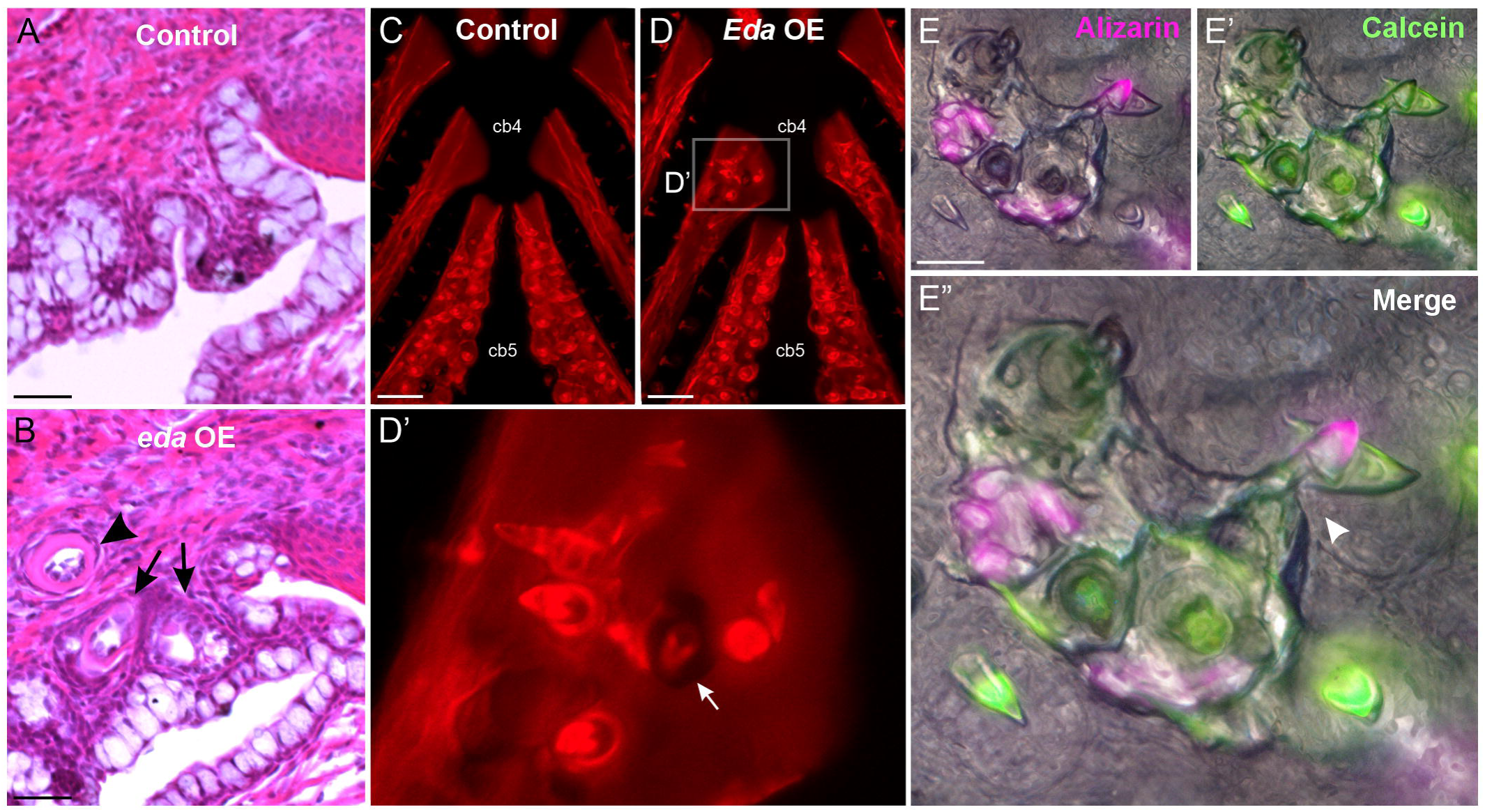
Ectopic pharyngeal teeth undergo regeneration and replacement for months following discontinued *Eda* overexpression. (A,B) Hematoxylin and eosin-stained sagittal sections (anterior to left) from control (A) and *eda* OE (B) zebrafish at 2 months of age following three embryonic heat shocks. Tooth germs are indicated with arrows, a later stage tooth is indicated by an arrowhead. (C-D) Alizarin Red-stained pharyngeal skeletons from control (C) and *Eda* OE (D) sticklebacks 2.5 months after six heat shocks from 6-11 dpf. Anterior to top. Arrow in D’ indicates an ectopic tooth germ with evidence of active bone remodeling. (E) a live bone staining pulse-chase assay was used to observe new tooth production and tooth retention following embryonic *Eda* OE. E shows Alizarin Red, E’ shows calcein, E” shows an overlay. Teeth with any Alizarin signal contain retained bone and were thus present at the pulse (“retained”). Teeth with only calcein signal contain only newly formed bone and thus were not present at the time of the pulse, but only the chase (“new”). Arrowhead in E” indicates an active tooth replacement event. Scale bars in A,B=10 μm; C,D=200 μm; E=100 μm.

We next tested for similar self-renewal capabilities of stickleback ectopic pharyngeal teeth. First, we overexpressed *Eda* in sticklebacks once per day from 6-11 dpf and allowed a 2.5 month recovery period prior to analysis of skeletal morphology by Alizarin Red (Fig. 7C,D). We found that n=11/13 *Eda* OE fish had ectopic teeth in at least one location within their pharynx, with most fish (n=9/13) demonstrating four or more distinct ectopic tooth fields. In each case, evidence of ongoing bone remodeling was present, typified by thin or punctured bone present on the skeletal element underlying unerupted tooth germs (Fig. 7D’, arrow). Thereafter we performed a follow-up experiment where we overexpressed *Eda* at 5 dpf with a single heat shock and allowed the fish to recover until 4 months of age, at which point we used pulse-chase live bone staining to assay tooth turnover (a 24 hour pulse of Alizarin red and 16 hour calcein chase 18 days later, see methods). We found that all present ectopic pharyngeal tooth fields were actively undergoing tooth turnover, as evidenced by the presence of alizarin-negative, calcein-positive teeth in 5/5 fish with ectopic tooth fields (Fig. 7E), in some cases with clear signs of predecessor tooth resorption with an adjacent new tooth germ (Fig. 7E’’, arrowhead).

We next asked whether face teeth in sticklebacks were capable of being maintained or renewed. We subjected fish to a 2x daily *Eda* OE treatment from 17-27 dpf (20 total heat shocks), keeping only those fish that grew at least one face tooth for later sampling, as evidenced by *dlx2b* reporter expression at 28 dpf. Subsets of fish were thereafter sacrificed and stained with Alizarin Red at different timepoints. At least one face tooth was observed in the following fractions of fish at the specified recovery timepoints: 6 days, 92% (n=11/12); 12 days, 50% (n=4/8); 18 days, 8% (n=1/13). The single ectopic tooth observed at the final collection point was ankylosed to the dentary and no longer demonstrated GFP fluorescence from *dlx2b*:*eGFP*. Throughout all collection points, we observed no clear morphological evidence of tooth replacement, nor any bell-stage *dlx2b* reporter-positive tooth germs at the 12 or 18 day collection points. Thus, *Eda*-induced ectopic pharyngeal teeth, but not face teeth, appear capable of regeneration long after *Eda* OE treatments are discontinued.

### Expression domains of *Edar* and *Troy* correlate with tooth formation competency

We hypothesized that *Eda*-driven ectopic tooth formation competency would likely be coincident with the endogenous expression of one or more tumor necrosis factor receptors (TNFRs). Using hybridization chain reaction, we analyzed the expression of *Edar, Troy,* and *Relt*, three closely related TNFR genes all previously implicated in epithelial organogenesis (*Troy* is also known as *TNFRSF19* [XM_040167660.1] and *Relt* is also known as *TNFRSF19L* [XM_040183272.1]) (Charles et al., 2009; Harris et al., 2008; Kim et al., 2019; Kojima et al., 2000; Ohazama and Sharpe, 2004). We first assayed the expression of these three TNFR genes at 6, 8, and 12 dpf (Fig. S3). *Edar* and *Troy* were expressed in tooth germs as well as some naïve or non-dental cell types (detailed below), rendering them as plausible candidate genes for involvement with ectopic tooth formation. *Relt* expression, on the other hand, was detected in a highly specific manner only in ameloblasts (mature inner dental epithelium) at mid- to late-bell stage tooth germs. At 6 dpf, during the most sensitive interval for ectopic tooth formation at regions like hb1 and cb4, *Relt* was not detected anywhere in the ventral pharynx, rendering it a poor candidate for involvement with ectopic tooth specification.

We thus expanded our expression survey to capture 6, 10, and 14 dpf, and assayed the expression of *Edar* and *Troy* alongside *Eda* (Fig. 8). Notably, at 6dpf we observed *Troy*- and *Edar-*positive cell populations along the ventral pharynx, approximately coincident with the medial regions of hb1-3 and cb4 where ectopic teeth were observed following *Eda* OE (Fig. 8A,B, gray boxes). *Edar* and *Troy* were both expressed in all observed cap-stage tooth germs, including the pioneer teeth on cb5 (Fig. 8A’,A’’,B black arrowhead). *Edar* was expressed more robustly in bell-stage tooth germs compared to *Troy* (Fig. 8C’,C’’,D’,D’’, white and gray arrowheads). *Troy* expression was additionally detected in tastebuds at 10 and 14 dpf (Fig. 8C’’, white arrow), as well as gills and their presumed precursor domains (Fig. 8A’’,C’’, carets). *Edar* transcripts were also detected in gill rakers (Fig. 8D’, double black arrow). *Eda* expression was detected in mesenchyme in the tooth field, particularly in the cells surrounding tooth germs on the medial side of the cb5 tooth fields at 8+ dpf (Fig. 8A’’’,C’’’,D’’’, black arrows). *Eda* expression was also weakly detected in mesenchyme medial to cb4 and hb3 (Fig. 8C’’’,D’’’, brackets), as well as in epithelium flanking the gill rakers at 10 and 14 dpf (Fig. 8D’’’, double white arrow). At higher resolution, *Eda*, *Edar* and *Troy* were detected in complex dynamic patterns in both dental epithelia and mesenchyme (Fig. S5).

**Figure 8.**
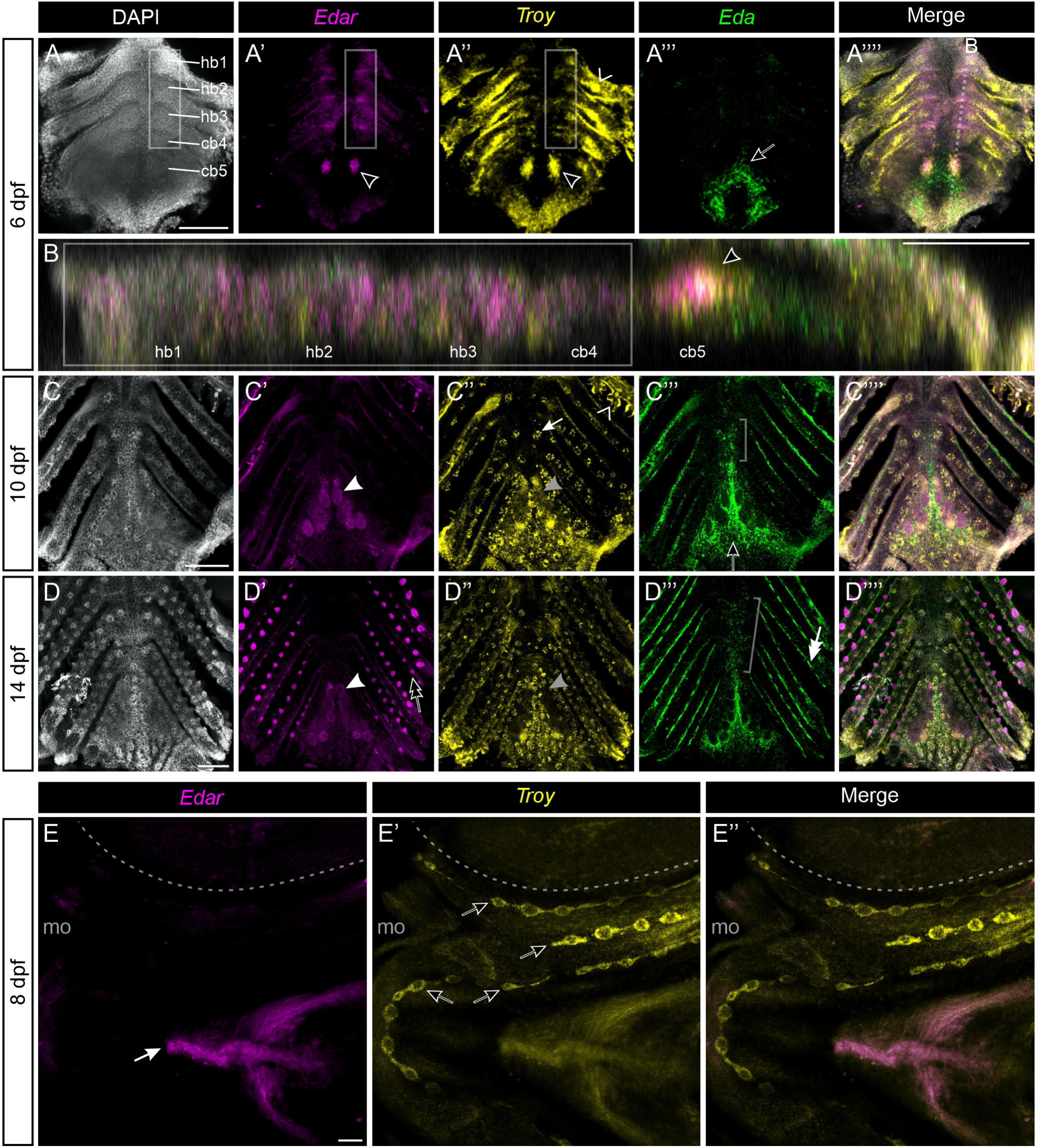
*Edar*, *Troy*, and *Eda* expression in the stickleback ventral pharynx and face. (A-D) Ventral pharyngeal preps of WT sticklebacks showing *Edar*, *Troy*, and *Eda* expression at 6, 10, and 14 dpf. Anterior to top in all panels, save B, where anterior is to the left. A,C,D show maximum intensity projections of Z-stacked optical slices, B shows a sagittal plane orthogonal reconstruction as indicated in A’’’’. Gray boxes indicate the competency region along the ventral pharynx. Black arrowheads in A’, A’’,B mark the right-side pioneer tooth on cb5. White arrowheads in C’,D’ mark bell-stage tooth germs. Double black arrow in D’ indicates a gill raker. Carets in A’’,C’’ indicate gill precursors and differentiating gills, respectively. Gray arrowheads in C’’,D’’ mark bell-stage tooth germs with little detectable expression. White arrow in C’’ marks a taste bud. Black arrows in A’’’,C’’’,D’’’ indicate medial mesenchyme at cb5. Brackets in C’’’,D’’’ indicate medial mesenchyme near hb3 and cb4. (E) *Edar* and *Troy* expression in a flattened facial prep at 8 dpf. Anterior to left, the eye is demarcated with a dashed line and the mouth opening is labeled “mo”. White arrow in E indicates ventral pharyngeal muscles. Black arrows in E’ indicate neuromast lines. Scale bars in A,C,D,E=100 μm, Scale bar in B=50 μm.

To test whether *Edar*, *Troy*, or *Relt* were expressed in a manner that might presage face teeth, we again tested for the expression of these three TNFRs at 8 dpf, representing the midpoint of an *Eda* OE treatment whereby we found >50% of fish formed facial teeth in the infraorbital region. *Relt* expression was not detected on the portion of the face we observed. *Edar* expression was observed in ventral pharyngeal muscles but was not detected in any region where facial teeth generally form (Fig. 8E). *Troy*, on the other hand, was found to mark neuromast primordia as well as interneuromast cells (Fig. 8E’). Thus, *Troy* but not *Edar* or *Relt* expression was detected in the external facial regions where teeth are competent to form.

## Discussion

### Ectopic tooth specification via *Eda* overexpression is most efficient during pioneer tooth formation stages

Using different sets of heat shock treatments to induce *Eda* expression, we established induction competencies through time for different regions capable of forming ectopic teeth in both sticklebacks and zebrafish. In both species, ectopic pharyngeal teeth can be activated via a lower number of *Eda* doses during stages where endogenous tooth field activation is taking place compared to earlier or later stages. However, in sticklebacks, the competency to form teeth at some of these locations is maintained through late larval stages, though more *Eda* OE doses were required, which still yielded relatively low tooth formation rates. We thus infer that the regions of the head skeleton that retain this latent developmental potential to form teeth undergo a short period during which they are most receptive to tooth induction via *Eda*, thereafter becoming more recalcitrant to tooth induction. Experiments using *Eda* mutants combined with *Eda* OE could resolve whether or not endogenous tooth fields follow a similar “competency arc” whereby their disposition to tooth induction changes through ontogeny. If endogenous tooth fields do exhibit temporal changes in competency through time, this could have implications for human health applications: perhaps *in utero* treatment with Fc-EDA during pioneer tooth initiation stages at 6-8 weeks would restore a greater proportion of the dentition than did the treatment at 26+ weeks, which was more efficient at restoring sweat glands (Schneider et al., 2018), which initiate at 20+ weeks of gestation.

One unexpected detail revealed by *Eda* ISH in sticklebacks is that *Eda* expression in wild-type sticklebacks is present along the posterior midline of the pharynx, medial to the cb4 and hb3 competency domains (Fig. 8C’’’,D’’’, brackets). This endogenous expression domain may partly explain why cb4 is particularly inclined to undergo tooth formation, as there is already a baseline level of *Eda* present in this region, lowering the amount of exogenous *Eda* required to elicit the tooth formation response. However, the generally less competent hb3 also appears to be subject to this expression domain, challenging this explanation.

### Pharyngeal tooth fields, once activated, can sustain themselves via regeneration

Previous work used an overexpression construct that could not be modulated by experimental intervention. Here we employed a heat shock promoter, allowing us to control the number and timing of *Eda* doses. By initiating teeth at early stages and then rearing fish for several months at normal temperatures, we were able to determine that ectopic pharyngeal tooth fields can persist by undergoing tooth regeneration. The teeth present at these later stages were thus themselves unlikely to have ever been exposed to exogenous *Eda* from the OE transgene but instead are utilizing a genetic program to trigger regeneration as part of the typical developmental trajectory of a tooth. The presence of tooth regeneration in these regions lends further support to these ectopic pharyngeal teeth as being otherwise typical, *bone fide* teeth. Interestingly, face teeth do not appear to have the capacity to replace themselves by engaging in tooth regeneration. Only 1/13 individuals with confirmed face tooth initiation maintained a tooth 18 days after the last heat shock, and this single tooth represented a rare case where it had become ankylosed to the underlying lower jaw bone (the dentary). There are two main possible explanations for this lack of regenerative potential: 1) ectopic face teeth are inherently different from ectopic pharyngeal teeth or endogenous teeth, and fail to specify successional epithelia and/or mesenchymal progenitors that are required for the maintenance of a tooth field, or 2) these face teeth can or even do specify the necessary cell types to engage in regeneration, but the general lack of ankylosis on the face leads to their relatively rapid loss due to typical epidermal sloughing or mechanical disturbances, causing the face teeth to simply fall off before they begin the process of regeneration. It is thus possible that the single retained face tooth we observed may have been poised to replace itself via regeneration. Further analyses of gene expression associated with successional dental cell types, and additional recovery experiments with longer observation windows could resolve these possibilities.

### Tooth organ specification

Outside of mammals, teeth are found at a litany of morphological locations beyond the maxilla and dentary bones. Reptiles, amphibians, and all major groups of fishes have member species that specify teeth on at least one bone of the head skeleton besides the dentary or maxilla (Berkovitz and Shellis, 2023a). Meanwhile, the early morphogenesis of individual tooth organs is a staunchly conserved process across species, demonstrating strikingly similar gene expression profiles to specify the cell types (odontoblasts and ameloblasts) that produce the characteristic specialized bony tissues found in teeth (dentine and enamel and/or enameloid) (Berkovitz and Shellis, 2023b). Given that individual teeth are widely accepted as homologous organ units (Sire et al., 2009), it is apparent that the relocalization of tooth initiation is not only developmentally feasible, but has occurred often. Ray finned fishes in particular demonstrate diverse patterns of tooth distribution. Besides being diverse overall, tooth distributions in this group have also been found to be highly labile in evolution: previous work provided strong support for widespread and parallel gain, loss, and even parallel regain following loss of certain tooth fields (Jandzik and Stock, 2021). This lability lends support to a model where, at least in some species, regions of the head skeleton are predisposed to be competent to execute a tooth formation genetic program, having maintained a “latent developmental potential” over the course of tens or even hundreds of millions of years of evolution (Aigler et al., 2014). Such developmental potential could ostensibly require only one or a few mutations to reactivate. For example, we would predict that gene regulatory changes that cause an expansion of *Eda* expression domains should be sufficient to reactivate tooth fields in some species, including sticklebacks and zebrafish.

At least three experimental conditions have been previously shown to increase primary tooth number or cause ectopic teeth to form: exogenous *Eda* and Retinoic acid have been shown to cause ectopic tooth fields in fish models, while *Osr2* loss-of-function has been shown to expand the endogenous molar tooth field in mice. *Eda* particularly presents a strong case for potential involvement in dental rearrangements over evolutionary time because: 1) Homo- or hemizygous null mutations in *Eda* have never been reported to cause inviable developmental defects in any species to our knowledge, 2) widespread overexpression of *Eda* elicits localized responses that prompt ectopic and possibly atavistic tooth fields to form in a limited set of morphological locations, 3) maternal delivery of exogenous Eda to pregnant mice or humans can rescue *Eda* mutant phenotypes in offspring without causing adverse developmental defects, and 4) most genetic *Eda* OE treatments yield viable fish with altered dental morphology, unlike published RA treatments or *Osr2* loss-of-function mutations, which are typically lethal (Gibert et al., 2019; Zhang et al., 2009). Thus, *Eda* seems to generally be tolerated in excess, or as a null allele, suggesting that most mutations that affect *Eda* expression would at least not be lethal, making *Eda* genes excellent targets for selectable mutations. Of course, it remains possible that more subtle variation in RA signaling titer or slight modulation of the *Osr2* regulatory network may work to achieve additional teeth in a viable manner. Experimental modulation of these and other pathways thought to be involved with epithelial appendage localization will shed light on whether other pathways similarly influence the initiation of teeth or other epithelial appendages.

### *Eda* genes and dental evolution

As previously hypothesized, here we provide additional evidence that *Eda* was duplicated at the Teleost Whole-Genome Duplication (TGD), creating type A and B paralogs, followed by differential retention and loss of *Eda* paralogs in various teleost groups. Here we show that stickleback fish subjected to overexpression of their own *Eda* (type A) are also capable of prompting ectopic tooth development, notably in many regions where other related fish species specify teeth. For example, some bass species (Beloniformes), which as Neoteleosts like sticklebacks we infer possess only type A *Eda*, are known to demonstrate teeth on hb2, hb3, and the bh (Gosline, 1985), all of which are regions where sticklebacks are capable of specifying ectopic teeth under *Eda* OE conditions. Thus, both type A and type B Teleost *Eda* duplicates, which diverged around 350 million years ago during early Teleost evolution, have retained the capability to prompt ectopic teeth. Given the general lability of the distribution of tooth fields across all teleosts, including those species with exclusively type A or type B *Eda*, we speculate that both teleost *Eda* paralogs have contributed to the evolvability of tooth distribution in the pharynx of teleosts.

Contrary to all observed ectopic pharyngeal tooth fields, the face teeth observed in sticklebacks do not correspond to teeth widely observed in other teleosts. Notably, there is at least one species that does exhibit such teeth: the denticle herring (*Denticeps clupeoides*). This species indeed specifies small teeth (denticles) on its infraorbitals, though notably it also adorns a far greater number of regions of its face with additional teeth, including the braincase and branchiostegal rays, where we never observed ectopic teeth (Greenwood and Greenwood, 1968). Taking a broader view, most Chondrichthyan species adorn their exteriors with denticles (Nicklin et al., 2024), leaving open the possibility that their natural presence in *Denticeps* and their experimental appearance in sticklebacks could be a harkening back to a rarely-utilized, but persistent developmental competency. *Eda* OE experiments in additional fish species will help resolve whether this competency is present beyond a select few species. On the other hand, it remains possible that face teeth in sticklebacks represent a fundamentally different kind of cellular transformation than is observed in the pharynx: rather than activating a latent developmental potential to form *bona fide* tooth fields, perhaps what occurs on the face realizes only enough of the tooth development program to form individual teeth that otherwise lack some critical surrounding components that comprise a true dental arcade. This alternative is supported by the apparent inability of face teeth to be maintained, unlike the pharyngeal teeth, and the rare occurrence of any bone of attachment. With this in mind, we speculate that the developmental path employed during ectopic pharyngeal vs face tooth initiation may not necessarily use the same pathways or downstream genetic mechanisms, despite their common entry point (*Eda* activation). Further experiments with various TNFR mutants will help resolve the pathway used to activate ectopic teeth in the pharynx and on the face.

### Expression patterns of TNFR genes give insights into potential receptor utilization during ectopic tooth formation

We performed expression analyses of the gene encoding the canonical Eda receptor, *Edar*, and two of its most closely related receptor genes, *Troy* and *Relt*, to determine which of these TNFRs, if any, are expressed in a manner that presages ectopic teeth. These analyses revealed that *Relt* expression was detected in a strikingly specific manner in mature ameloblasts and is thus a poor candidate gene with respect to what confers the ectopic tooth formation response to Eda. On the other hand, *Edar* and *Troy* expression both mark regions of the ventral pharynx where ectopic teeth are capable of forming, making them excellent candidates for involvement with conferring ectopic tooth formation in the pharynx. In the infraorbital and mandibular regions were face teeth most typically form, we found only *Troy* expression in nascent neuromasts and interneuromast cells. Given our observation that face tooth formation and neuromast counts are anticorrelated, we speculate that Eda can signal through Troy, and that this interaction can cause cells in the neuromast lineage to take on a fate of the dental epithelium. Notably, neuromasts are not present inside the pharyngeal cavity, thus ectopic pharyngeal tooth epithelium is highly unlikely to follow this same developmental route. Further tests using *Troy* mutants and *Eda* overexpression could test whether this receptor is necessary for conferring any of the effects observed during *Eda* overexpression. These and other experiments could additionally test whether the molecular basis of competence for organ formation in this system is defined by differential, or perhaps combinatorial, receptor expression.

## Conclusion

Here we show that the competency to respond to ectopic *Eda* by initiating tooth organs shifts anatomically within the head and temporally across development. In both zebrafish and sticklebacks, ectopic pharyngeal tooth fields are capable of maintaining themselves through tooth replacement in the absence of additional heat shock-driven *Eda* expression, suggesting these teeth can act in a semi-autonomous fashion to renew themselves once initiated. However, face teeth in sticklebacks do not appear to regularly engage in replacement and instead are usually lost within 18 days. Finally, expression analyses of three TNFR genes in sticklebacks revealed that *Edar* and *Troy*, but not *Relt*, are expressed in a manner consistent with a potential role in conferring the ectopic tooth initiation response. Together these data reveal that during development, the competency to respond to a secreted signal and initiate organogenesis is surprisingly dynamic and suggest that differential receptor expression could be a molecular mechanism underlying this dynamic competency.

## Methods

### Amino acid alignment and phylogenetic analysis

Eda amino acid sequences were collected from the National Center for Biotechnology Information (NCBI) GenBank and Ensembl. Accession numbers for each sequence are listed in Supplementary File 1. Genomicus versions 99-106 were additionally used to browse synteny data and predicted orthology relationships (Nguyen et al., 2022, 2018). Animo acid sequences were aligned with MUSCLE (Edgar, 2004) in MEGA7 (Kumar et al., 2016). The tree was inferred by using the Maximum Likelihood method and Whelan and Goldman (WAG) model (Whelan and Goldman, 2001) in MEGA11 (Tamura et al., 2021). The tree with the highest log likelihood (−10634.31) is shown in Fig. S1. Initial tree(s) for the heuristic search were obtained by applying the Neighbor-Joining method to a matrix of pairwise distances estimated using the Jones-Taylor-Thornton (JTT) model (Jones et al., 1992). This analysis was performed on 37 amino acid sequences from 26 species. All positions in the alignment with less than 80% site coverage were eliminated, leaving a total of 337 positions used to build the tree. A bootstrap analysis was conducted with 100 replicates; the percentage of trees in which the associated taxa clustered together is shown near each node in Fig. S1. The resulting tree was rearranged and some formatting was performed in FigTree version 1.3.1 (Rambaut and Drummond, 2009).

### Transgene plasmids

Plasmids for transgenesis were created as previously described (Square et al., 2023). The zebrafish *eda* coding sequence (NCBI reference sequence NM_001115065.1) with an antisense T3 site after the stop codon on the 3’ end was synthesized by Gene Universal (Delaware, USA) and restriction cloned into the pT2overCherry construct using XbaI and XhoI. Stickleback *Eda* was codon optimized and synthesized by Integrated DNA Technologies (Iowa, USA). The codon optimized gene was designed to output a protein product that matches NCBI reference sequence XP_040030617.1. Codon optimized stickleback *Eda* was then restriction cloned into pT2overCherry using XbaI and XhoI.

The 4122 bp zebrafish *dlx2b* promoter and upstream regulatory region was amplified from zebrafish genomic DNA using the primers 5’- GAGTCATTTTGATCTGGAGAAAGCTGATG-3’ and 5’-TTCGCAGGAAGAAGAGACTACTCAACG-3’. The product was reamplified with the primers 5’-gccgggatccGAGTCATTTTGATCTGGAGAAAGCTGATG-3’ and 5’- ggtggtgtcgacCGCAGGAAGAAGAGACTACTCAACG-3’ to add BamHI and SalI to the 5’ and 3’ ends of the product, respectively. The reamplified product was digested with BamHI and SalI. The pT2He plasmid (Howes et al., 2017) was digested with BamHI and SalI in parallel, and the smaller insert containing the heat shock promoter was discarded. The digested PCR product and remainder of the pT2He plasmid backbone (without the hsp70l promoter) were ligated together, creating pT2_Dr_d2b, which drives eGFP expression directly under the control of the 4122 bp zebrafish promoter and upstream regulatory region, recreating the main components of the previously published transgene cassette (Jackman and Stock, 2006). Once sequence verified with Sanger traces from each end of the new inserts, plasmids were midiprepped (QIAGEN), phenol-chloroform extracted, precipitated, and resuspended in DEPC-treated water per standard methods.

### Fish husbandry and transgenic line establishment

All husbandry and experiments were performed with approval of the Institutional Animal Care and Use Committee of the University Florida (protocol IACUC202300000692) or the University of California-Berkeley (protocol AUP-2015-01-7117). Sticklebacks were raised in 110 liter aquaria at 18°C in 3.5 g/L Instant Ocean salt and 39.4 mg/L sodium bicarbonate. Sticklebacks (Cerrito Creek [CERC]) were fed a common diet of live Artemia as young fry, live Artemia and frozen Daphnia as juveniles and frozen bloodworms and Mysis shrimp as subadults and adults. Zebrafish (AB strain) were raised according to standard methods (Sprague et al., 2001). At the end of each experiment, fish were euthanized with MS-222. Reporter transgene fluorophores and DASPEI stained-fish were imaged without fixation or alcohol exposure. Alizarin Red skeletal stains, pulse-chase labeling, and ISH material were then fixed in 4% formaldehyde overnight at 4°C or for 4h at RT. Fixative was washed out with one rinse and 2x 10 min PBST washes. Live pulse-chase bone labeling was imaged without alcohol exposure, all other assays were followed by rinsing and storage in MeOH (*in situ* material) or EtOH (skeletal stains or H&E).

Transgenesis was accomplished in sticklebacks and zebrafish by coinjecting Tol2 mRNA and prepared plasmids described above. In the case of the *Eda* OE and *dlx2b* reporter transgenes in sticklebacks, fish were injected (Erickson et al., 2016), outcrossed, and screened as previously described (Square et al., 2023) to achieve and maintain a single insertion transgenic background for all experiments herein. The zebrafish *eda* OE line (ZFIN accession bk407tg) was uniquely and intentionally maintained as multi-insertion in order to increase the heat shock dosage due to the generally weaker effects of this transgene. For the experiments shown in Fig. 7C, all assessed individuals resulted from outcrosses of the same male fish, who we infer carried three insertions of the transgene (201/237 mCherry+, ∼84.8%).

### Heat shock transgene activation

Heat shocks were administered essentially as previously described, with some modification (Square et al., 2023). Zebrafish were confined to a 50 mL tube in 25 mL of embryo medium and placed in a 38°C water bath for 65 minutes (it takes ∼5 minutes for 25 mL of fish water to ramp ∼10°C). For sticklebacks, single heat shocks prior to 10 dpf were performed as described for zebrafish using 50 mL tubes and a water bath. For multiple heat shocks in sticklebacks, or for heat shocks occurring between 10 and 30 dpf, stickleback fry were housed in a 2 L tank with light aeration and one 50 W aquarium heater set to 29°C, which would be powered on in two or more 2 hour intervals (it takes ∼50 minutes for 2 L of fish water to ramp ∼10°C). When necessary, fish were lightly anesthetized with MS-222 and sorted based on their fluorescence profile. “Control” fish refer to heat-shocked siblings that did not inherit the OE transgene and experienced the same heat shock regiment as their OE transgene-carrying siblings, except those shown or described in Fig. 1D “Control (HS-)”, Fig. 5A “0 shocks,” and Fig. 6B “0 shocks” which indicate OE transgene carriers that were not heat shocked.

### Alizarin Red skeletal staining

Alizarin Red S was used for skeletal staining essentially as previously described (Ellis and Miller, 2016). Samples were stained in a 0.008% Alizarin Red S solution in 1% KOH for 24+ hours, replacing the solution if necessary (if the solution turns pale). Large samples from subadults were sometimes left in Alizarin Red staining solution for up to one week. Once adequately stained, samples were transferred to 0.5% or 1% KOH for 1-3 days to continue clearing, if necessary. Tooth plates were then dissected and scored and/or imaged by further clearing in 50% glycerol for 1-5 days before flat mounting in 50% or 90% glycerol in 1x PBS.

### Live pulse-chase bone labeling

Live pulse-chase bone labeling on sticklebacks was performed essentially as previously described (Square et al., 2023). Following heat shock treatments and long recovery windows as described above, fish were placed into a tank containing an Alizarin Red live staining solution (0.1 g/L Alizarin Red S with 1 mM HEPES) made in otherwise normal tank water (Ellis et al., 2015). Sticklebacks were pulsed with Alizarin Red for 24h in 2L of Alizarin Red live staining solution. Fish were then rinsed 2x and washed 2x 20 minutes in normal tank water before returning to their standard 110L tank in the vivarium. 18 days later, fish were chased with Calcein live staining solution (0.05 g/L calcein with 1 mM sodium phosphate in normal tank water) for 16h in 2L. Fish were then rinsed 2x and washed 4x 20 minutes in normal tank water before being euthanized and fixed. The next day, samples were rinsed 1x and washed 2x 20 min in tap water, and pharyngeal skeletons were dissected out and treated with 1% KOH overnight at RT (without a storage/EtOH step). Skeletons were then rinsed in tap water and washed 2x 5 min in PBST and prepared for mounting and imaging by stepping the samples through 30, 60, and 90% glycerol in 1x PBS. Samples were then flat-mounted and imaged essentially as previously described (Ellis and Miller, 2016; Square et al., 2023).

### *In situ* hybridization

To visualize mRNA distributions, *in situ* hybridization (ISH) was carried out either by a traditional wholemount colorimetric method or Hybridization Chain Reaction (HCR). Colorimetric wholemount ISHs were performed essentially as previously described (Square et al., 2021). The stickleback *Pitx2* riboprobe was previously published (Ellis et al., 2016; Square et al., 2021). HCR was carried out using proprietary probe sets developed by Molecular Instruments (California, USA) and by following a modified version of the manufacturer’s protocol, namely using a higher concentration and volume of probe and probe hybridization solution (16 pmol in 1.8 mL), and material fixation/preparation was performed as outlined above.

### DASPEI staining

DASPEI (2-(4-(Dimethylamino)styryl)-N-ethylpyridinium iodide) was used to visualize neuromast position on live stickleback fry (Wark and Peichel, 2010). Sticklebacks were anesthetized and live stained with 0.02% DASPEI in normal tank water plus MS-222 for 5 minutes before washing in normal tank water plus MS-222 and imaging.

### Sectioning and H&E staining

Hematoxylin and Eosin staining was used to observe ectopic teeth in zebrafish. Following fixation, zebrafish heads were sectioned on the sagittal plane at a thickness of 7 um. Slides were subjected to a series of staining and destaining washes, coverslipped, and imaged as described previously (Square et al., 2021).

## Supporting information

Supplementary File 1 (accession numbers)

## Acknowledgements

We thank David Stock, David Jandzik, Sophie Archambeault, Mark Stepaniak and Megan Martik for input on experiments and feedback on the manuscript; Tyler Mentley, Frances Campbell, and Kait Kliman for assistance with zebrafish husbandry.

## Author contributions

Conceptualization: T.A.S.; Methodology: T.A.S., C.T.M.; Software: T.A.S., Z.Z.C.; validation: T.A.S., Z.Z.C., S.N.N.; Formal analysis: T.A.S.; Investigation: T.A.S., E.J.M., S.S., Z.Z.C., S.N.N., L.M.S., P.Q.H.; Resources: T.A.S., C.T.M.; Writing - original draft: T.A.S.; Writing - review & editing: T.A.S., C.T.M.; Visualization: T.A.S.; Supervision: T.A.S., C.T.M.; Project administration: T.A.S., C.T.M.; Funding acquisition: T.A.S., C.T.M.

---

**Figure S1.**
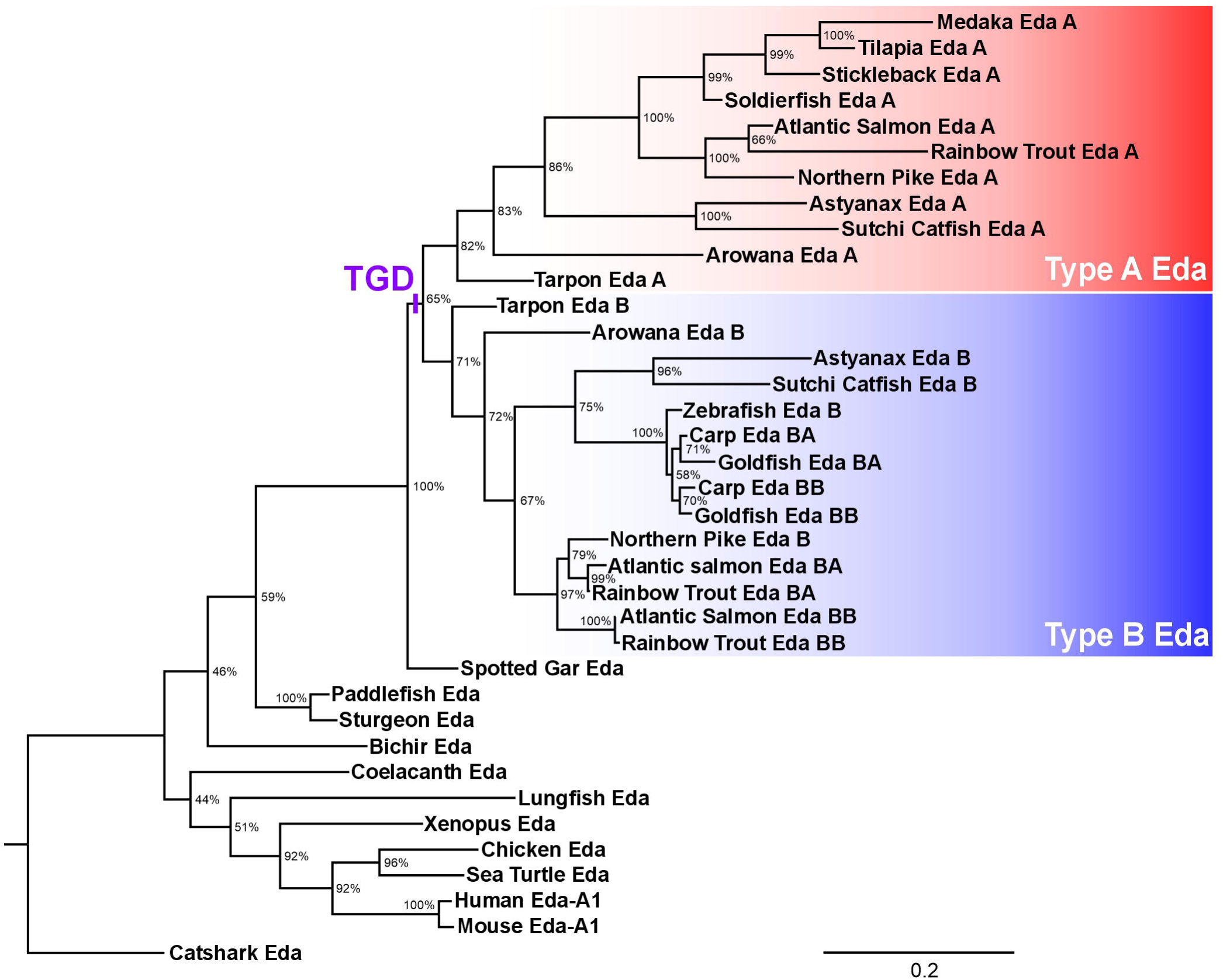
Phylogenetic tree of Eda amino acid sequences in jawed vertebrates. A Maximum Likelihood reconstruction based on an alignment of 37 Eda sequences from selected species. See Supplementary File 1 for a table of accession numbers for the sequences included. A 100 replicate bootstrap test was performed, all resulting values are shown near each node. Catshark Eda was selected as the outgroup. See methods for aa alignment and tree-building parameters. TGD, Teleost Genome Duplication.

**Figure S2.**
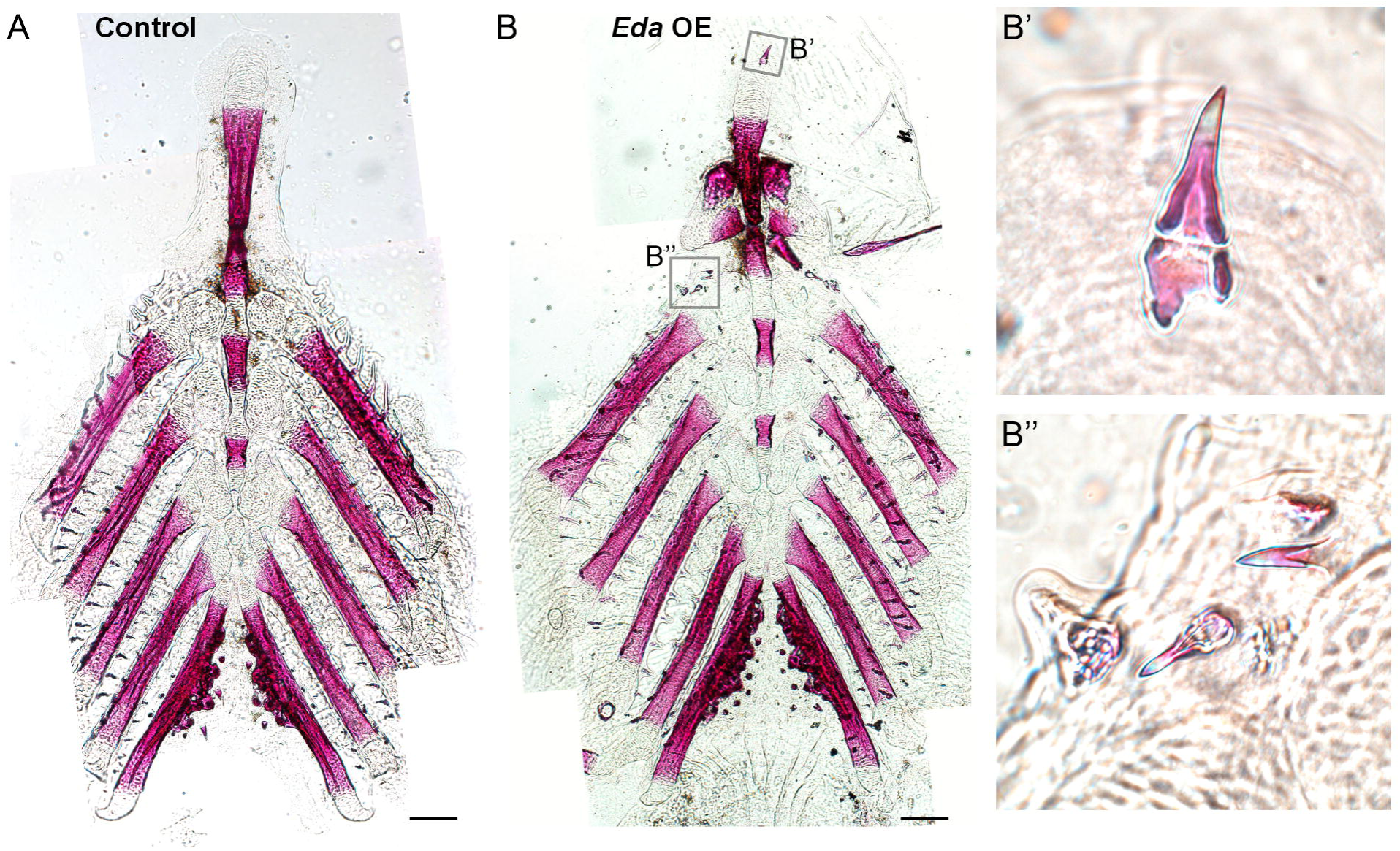
Ectopic pharyngeal teeth resulting from larval *Eda* overexpression. Alizarin Red-stained pharyngeal preps from control (A) and *Eda* OE (B) fish that underwent the larval OE treatment. Insets are indicated in the *Eda* OE treatment (B’ and B”) to show detail of ectopic teeth in the *Eda* OE condition. Scale bars=200 μm.

**Figure S3.**
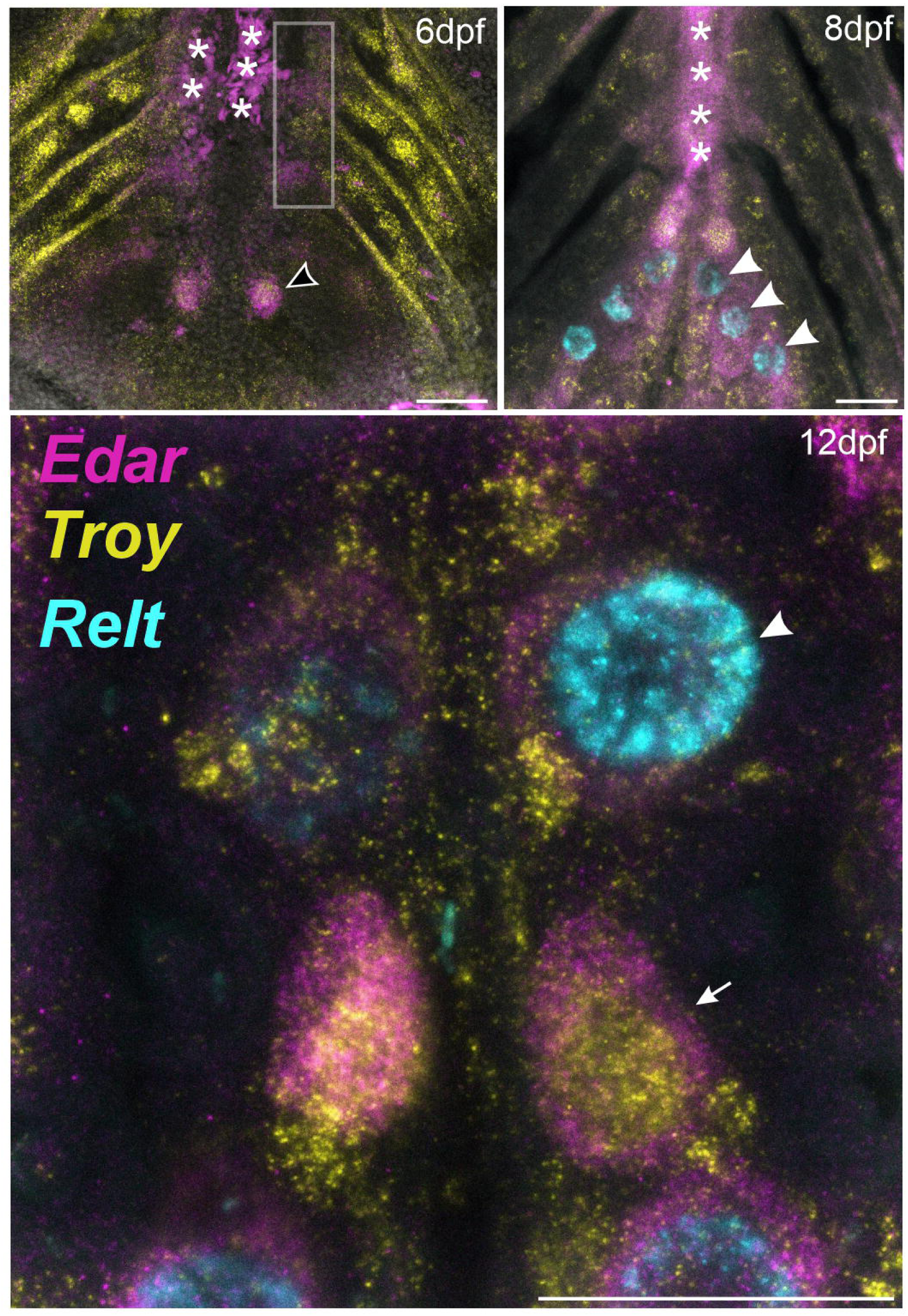
*Edar*, *Troy*, and *Relt* expression in the stickleback pharynx. *Edar* and *Troy* transcripts were detected in pioneer teeth at 6 dpf (black arrowhead) and non-dental tissues presaging tooth formation competency (gray box). *Relt* is not present at 6 dpf and was only detected in ameloblasts during tooth differentiation at 8 and 12 dpf (arrowheads). White arrow marks a cap-stage tooth germ at 12 dpf. White asterisks mark autofluorescence from blood cells trapped near the heart. Scale bars=50 μm.

**Figure S4.**
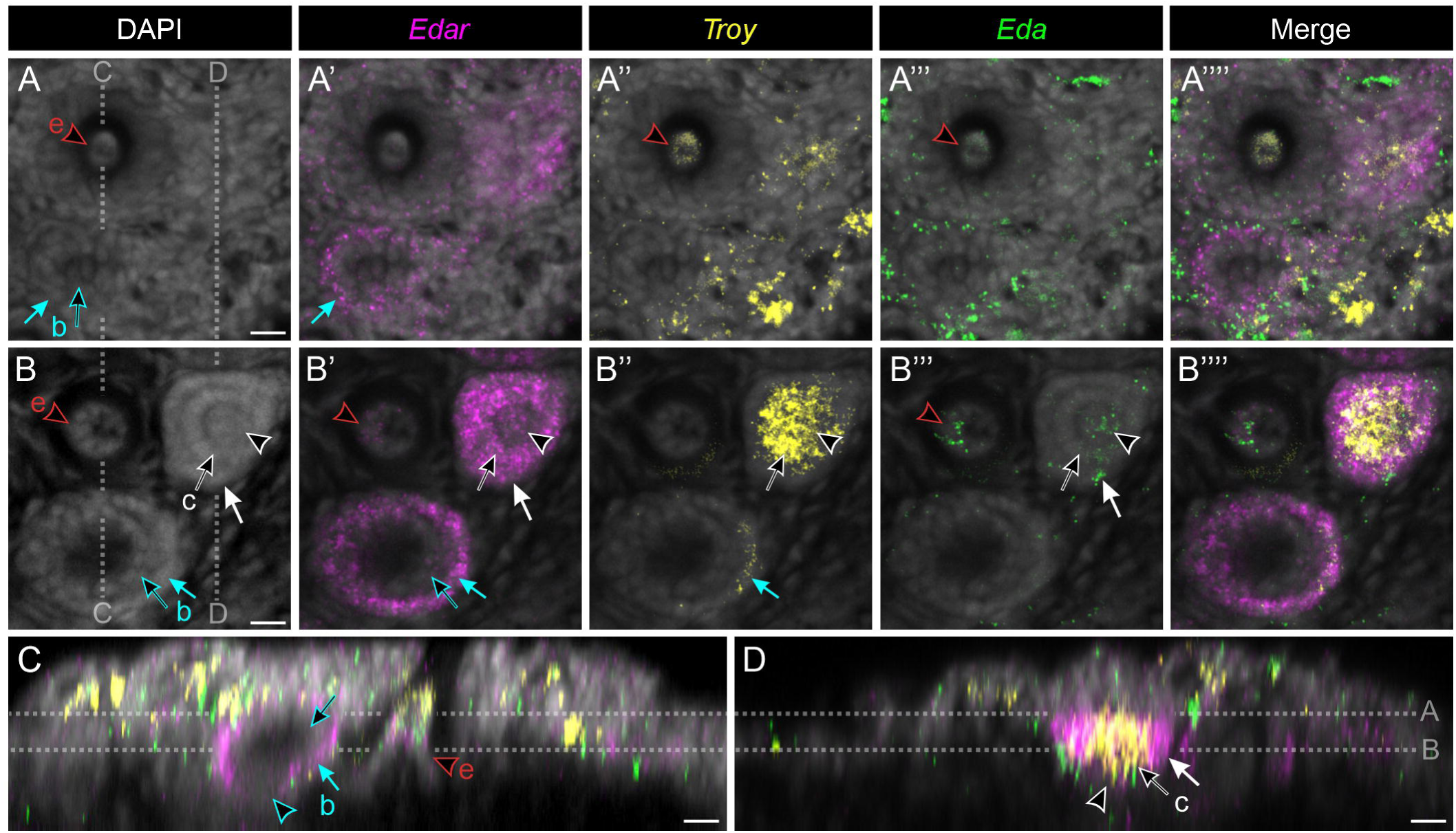
*Edar*, *Troy*, and *Eda* expression within and surrounding tooth germs. A cluster of three tooth organs on dorsal tooth plate 2 (DTP2) at 18 dpf are shown as example to further compare tooth germ expression at higher resolution. (A,B) optical sections showing a shallow (A) and deep (B) plane of section reveal dental mesenchyme of an erupted tooth (“e” in A-C, red and black arrowheads), bell-stage tooth germ (“b” in A-C) inner dental epithelium (black and cyan arrows) and outer dental epithelium (cyan arrows), and cap-stage (“c” in A,B,D) mesenchyme (black and white arrowhead), inner dental epithelium (black and white arrow), and outer dental epithelium (white arrow). All tissues are labeled in the DAPI panels, while the subsequent panels show arrows or arrowheads only if transcripts were detected in that tissue. Merged panels are shown without markup. Gray dotted lines in A and B labeled “C” and “D” indicate the plane of the orthogonal views in panels C and D. We detected *Edar, Troy,* and *Eda* within and/or surrounding tooth organs of most developmental stages. Cap stage tooth germs (“c”) demonstrated the most widespread expression of all three genes, where *Edar* was detected in both inner and outer dental epithelium (B’, black and white arrows) as well as dental mesenchyme (B’, black arrowhead), *Troy* most strongly marked inner dental epithelium (B’’, black arrow) and mesenchyme (B’’, black and white arrowhead), and *Eda* sparsely marked inner and outer dental epithelium and mesenchyme (B’’’, black arrowhead). Bell-stage tooth germs (“b”) demonstrated more restricted expression, with *Edar* mainly detected in the outer dental epithelium (B’, cyan arrow) with some limited *Troy* signal in the same tissue (B’’, cyan arrow), and little signal seen in mesenchyme (C, cyan and black arrowhead). Erupted tooth mesenchyme (“e”) expressed *Troy* apically (A’’, red and black arrowhead) while demonstrating limited *Edar* and *Eda* signal at a deeper level (B’,B’’’, black arrowheads). (C,D) Orthogonal composite views confirmed this general scheme. Labeling and arrow scheme same as above, all tissues are marked with arrows regardless of detected expression. Scale bars=10 μm.

